# Extensive genome introgression between domestic ferret and European polecat during population recovery in Great Britain

**DOI:** 10.1101/2021.12.20.473448

**Authors:** Graham J Etherington, Adam Ciezarek, Rebecca Shaw, Johan Michaux, Elizabeth Croose, Wilfried Haerty, Federica Di Palma

## Abstract

The European polecat (*Mustela putorius*) is a mammalian predator which occurs across much of Europe east to the Ural Mountains. In Great Britain, following years of persecution the European polecat has recently undergone a population increase due to legal protection and its range now overlaps that of feral domestic ferrets (*Mustela putorius furo*). During this range expansion, European polecats hybridised with feral domestic ferrets producing viable offspring. Here we carry out population-level whole genome sequencing on domestic ferrets, British European polecats, and European polecats from the European mainland and find high degrees of genome introgression in British polecats outside their previous stronghold, even in those individuals phenotyped as ‘pure’ polecats. We quantify this introgression and find introgressed genes under selection that may assist in cognitive function and sight.

## Introduction

Mustelidae form the largest family of the order Carnivora, comprising around 60 species. *Mustela*, a genus of Mustelidae that originated at least 5.3 MYA, contains around 17 species of weasels, stoats, mink, polecats, and ferrets (Koepfli et al., 2008). The European polecat (*Mustela putorius*) occurs in a range of habitats including farmland, wetland, and woodland, where it feeds mainly on rabbits, rodents, amphibians, and small birds (Croose, 2016; Lodé, 1997). The domestic ferret (*M. putorius furo*) has long been associated with man and is thought to have been domesticated around 2000 years ago and introduced in Great Britain around the 11th Century (Thomson, 1951). The ancestral species to the ferret is also disputed with most authors treating the domestic ferret as a descendant of the European polecat, but others treating it as a descendant of the Steppe polecat (*M. eversmanii*) (e.g. (Blandford & Walton, 1991; Driscoll, Macdonald, & O’Brien, 2009)).

In Great Britain, the European polecat (hereafter ‘polecat’) has a chequered history. Once widespread across most of the country, it was systematically persecuted to near-extinction in the late 1800s when it became confined mainly to small areas of central Wales and the Welsh border counties (Figure *1* a) (Croose, 2016; Langley & Yalden, 1977). Following legal protection and a reduction in persecution, polecat numbers started to increase from the 1930s, seeing a range expansion that now reaches almost every county in England and Wales along with limited areas in Scotland (Figure *1* b) (Croose, 2016).

**Figure 1.**
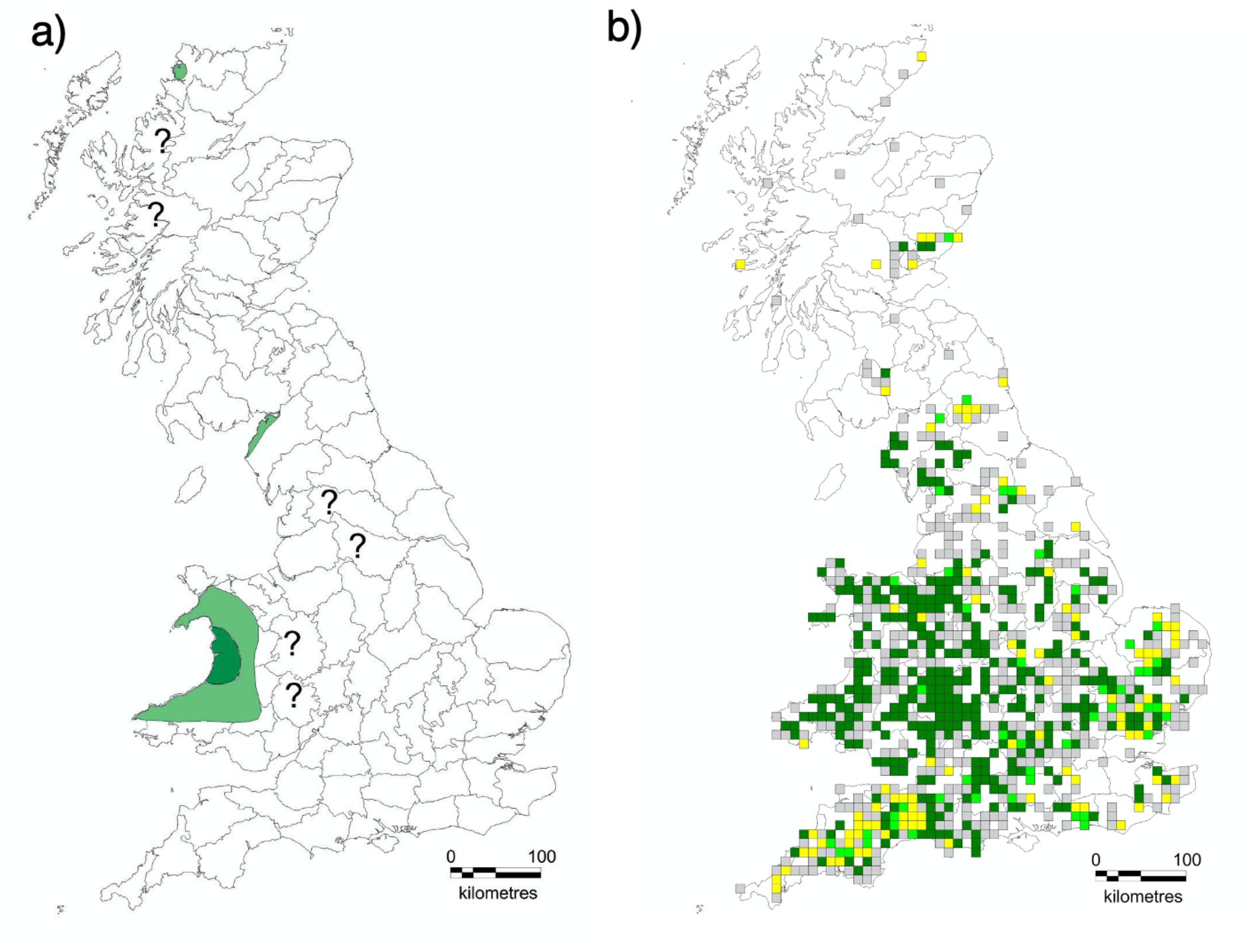
a) The range of the European polecat across the Great Britain in 1915 (dark green indicates stronghold, pale green indicates localised or rare occurrences, and ‘?’ indicates uncertain status. b) The distribution of hectads in which verifiable records of true polecats (dark green), polecat-ferrets (yellow), both true polecats and polecat-ferrets (lime green) and unverifiable records (grey) were received during the 2014-2015 European polecat survey (Croose, 2016).

During their range expansion, polecats hybridised with feral domestic ferrets most frequently at the edge of the polecats’ range (Costa et al., 2013; Croose, 2016). Previous studies focusing on polecat mitochondrial haplotypes reported between two (one ferret and one polecat (Davison et al., 1999)) and three haplotypes (one ferret and two polecat) (Costa et al., 2013) occurring in Great Britain. Further work using 11 microsatellite loci suggested genome introgression was quite prevalent, occurring in 31% of polecat samples. However due to an insufficient number of microsatellite loci, no F1 hybrids or backcrossed individuals could be identified (Costa et al., 2013). Polecats are difficult to phenotypically identify, with many individuals displaying a polecat phenotype but possessing ferret haplotypes, suggesting strong selection on the polecat phenotype (Birks & Kitchener, 1999; Costa et al., 2013).

Previous work on British polecats has concentrated mainly on the analyses of mitochondrial genes or used a small number of microsatellites that have lacked the power to quantify the extent of per-individual genome introgression. There are no studies that look at adaptive introgression in polecats and the amount of research relating polecat and ferret genotypes to phenotype is minimal. Here we carry out population-level whole genome sequencing on a range of domestic ferrets, polecats from the European mainland, and polecats and polecat-ferret hybrids from Great Britain and examine the extent of domestic ferret introgression in British polecats.

## Methods

### Sample origins

We sourced 45 *Mustela* samples, which comprised of 8 domestic ferret, 15 European polecats from the European mainland (2 from Spain, 3 from Austria, 3 from France, 5 from Italy, and 2 from Germany), 16 European polecats phenotyped as ‘pure’ and 3 phenotyped as polecat-ferret hybrids from Great Britain, 2 Steppe polecats (*M. eversmanii)*, and 1 Least weasel (*M. nivalis*) (Supplementary Table S1). The British samples were further sub-divided by location, with six samples from Wales (or bordering counties) and the remining 13 samples allocated to the ‘English’ group (pure and hybrids). All samples were sourced as tissue or DNA. Additionally, 4 Black-footed ferrets (*M. nigripes*) whole genome sequences were obtained from publicly-available data.

### Sample preparation and sequencing

DNA extraction was carried out using Qiagen DNeasy Blood and Tissue following the manufacture’s protocol. Samples were quality checked using the Nanodrop Spectrophotometer. Illumina short-read sequencing was carried out on all samples (Supplementary Table S2). The full protocol is described under ‘Library construction protocols’ at protocols.io under dx.doi.org/10.17504/protocols.io.bww9pfh6

The 8 domestic ferrets represent the most diverse set from 76 other ferret samples and were selected as follows. A Nimblegen capture array was designed, targeted to 2 Mb from each of the 6 largest scaffolds from the ferret genome assembly for a total of 12 Mb (Peng et al., 2014). All 76 samples used in the capture were drawn from multiple populations (US, China, Australia) and phenotypes (longhair, sable, albino, and cinnamon). The capture library was then sequenced on an Illumina HiSeq and GATK was then used to call SNPs (McKenna et al., 2010). Clustering and phylogenetic methods were then used to select 8 samples that represented the most diverse set from the original 76.

### Mitochondrial genome assembly

The program Mitoz ‘all’ (Meng, Li, Yang, & Liu, 2019) was used to filter the genomic reads for those of a mitochondrial origin, assemble them *de novo*, identify mitochondrial scaffolds and annotate each scaffold. Using the European polecat mitochondrial genome (accession NC_020638.1) as an anchor, assemblies were then re-ordered so they all started at the same relative positions. The full protocol is described under ‘Mitochondrial genome assembly’ at protocols.io under dx.doi.org/10.17504/protocols.io.bqzbmx2n

### Genomic variant calling and consensus assembly

Read quality was assessed using the FASTQC tool (Andrews, 2010). Reads with low quality scores (<Q20) were trimmed using the Trim Galore! tool (Krueger, 2015). Next, we assembled consensus genome sequences for each sample, following the Genome Analysis Toolkit (GATK, version 4.1.3) best-practice workflow for SNP and Indel calling in nonmodel organisms (Poplin et al., 2017), which is fully described under ‘GATK Nuclear variant discovery and consensus assembly’ at protocols.io under dx.doi.org/10.17504/protocols.io.bqzgmx3w

Briefly, reads were mapped using BWA MEM (Li & Durbin, 2009) to a new version of the domestic ferret reference genome MusPutFur1.0 scaffolded with Bionano data (accession GCA_920103865). After marking duplicates with Picard Tools (Broad_Institute, 2019), variants were called using the GATK HaplotypeCaller (using the PCR-free indel model where appropriate). The mean value of SNP quality, read depth, and mapping quality (QUAL, DP, and MQ values respectively from the VCF file) were calculated and GATK VariantFiltration was carried out marking SNPs with values lower than the mean scores as failing the filter. Using the variants that passed filtering, a recalibration table was created using GATK BaseRecalibrator and then Base Quality Score Recalibration was carried out using the table with the GATK ApplyBQSR tool to produce a recalibrated BAM file. This was then followed by a second round of GATK HaplotypeCaller using the recalibrated BAM file outputting a GVCF file (all sites, regardless of haplotype). Next, GATK GenomicsDBImport was used to import all GVCF files from all samples, followed by GATK GenotypeGVCFs to create a single multi-sample VCF file, followed by the creation of a SNP-only version using the GATK SelectVariants tool. Finally, consensus genome sequences were created for each sample using the GATK FastaAlternateReferenceMaker tool.

### Genome-wide SNP data

Whole genome SNPs for all 49 samples were filtered using PLINK v1.9 with an LD threshold of 0.8, within 5kb windows. (Chang et al., 2015). This subset of SNPs was further filtered to only include one alternate homozygous sample and one reference homozygous sample to alleviate any reference bias. SNPs that were not present across 3 samples were also filtered out. This resulted in 2,694,963 SNPs. These sites were then concatenated into phylip format using a custom python script for input into RAxML-NG.

### Phylogenetic analyses

#### RAxML

We used ClustalW to align the 49 *de novo* mitochondrial genome assemblies (Thompson, Gibson, & Higgins, 2002). We then used RAxML-NG to construct maximum likelihood (ML) phylogenies for the *de novo* mitochondrial genome sequences and the genome-wide SNP data (Kozlov, Darriba, Flouri, Morel, & Stamatakis, 2019). Full details are described under Step in ‘Population Structure Phylogenetics’ at protocols.io under dx.doi.org/10.17504/protocols.io.bretm3en. Briefly, we ran RAxML in two stages. The first is the ‘check’ and ‘parse’ options of RAxML. ‘check’ ensures for alignment correctness and removes empty columns. ‘parse’ creates a binary MSA file (named *raxml.rba). It also estimates the resources (cores and memory) needed to run RAxML efficiently. Using the resources advised by the previous step, we ran raxml ‘all’ which runs a ML tree search and non-parametric bootstrap by re-sampling alignment columns and re-inferring a tree for each bootstrap replicate.

#### TreeMix

We also used TreeMix to examine historical relationships among populations (Pickrell & Pritchard, 2012). TreeMix uses allele frequency datasets to estimate relationships among populations, using a graph representation that allows both population splits and migration events. Using the LD-pruned SNPs described above, we ran TreeMix under models allowing between 1 and 10 migration edges. Full details are described under Step 2 in ‘Population Structure and Phylogenetics’ at protocols.io under dx.doi.org/10.17504/protocols.io.bretm3en. To estimate the optimal number of migration edges, we used the R package OptM using the default Evanno method (Fitak, 2019). The number of migration edges was chosen based on a plateau of log likelihoods and when greater than 99.8% of the variance was explained.

#### Population Structure

We used model-based clustering implemented in ADMIXTURE to visualise the genetic ancestry of domestic ferrets and European polecats. After selecting samples with GATK SelectVariants, we used VCFTOOLS and Plink to reformat our data into the BED format required by ADMIXTURE. We ran ADMIXTURE with bootstrapping in order to estimate the most likely value of ‘K’. Full details are described under Step 3 in ‘Population Structure and Phylogenetics’ at protocols.io under dx.doi.org/10.17504/protocols.io.bretm3en

### Introgression

#### HyDe analysis

The consensus genome sequences for each sample were concatenated into a single fasta sequence, which were then in turn concatenated to construct a single multiple sequence alignment containing all samples. Variant sites were then extracted using SNP-sites (Page et al., 2016). We partitioned the data into 5 populations – domestic ferrets, mainland European polecats, Welsh polecats, English polecats, and weasel (outgroup). Phenotypic hybrids were not treated as a separate case in order to make no prior assumptions on genetic backgrounds assigned only by phenotype (which is notoriously difficult to assign correctly). HyDe was used to run the ‘full’ analysis (using HyDe’s ‘run_hyde.py’ script), which tests all combinations of three populations in all directions for evidence of hybridisation (with the addition of the weasel outgroup in each test). Hyde filters the results from the hybridization detection analysis to only include trios with significant results and sensible values. The filtered list of trios was then used as input to test hybridisation of individuals within the populations that have significant levels of hybridisation. Full details are described under Step 1 of ‘Introgression’ at protocols.io under dx.doi.org/10.17504/protocols.io.bq7tmznn

#### Dsuite analysis

ABBA-BABA tests (also known as Patterson’s D statistic) are commonly used to assess evidence of gene flow by examining patterns of allele sharing between populations or species in genomic datasets (Durand, Patterson, Reich, & Slatkin, 2011). Dsuite is a software package that implements ABBA-BABA tests between designated populations, as well as a number of other closely related methods (Malinsky, Matschiner, & Svardal, 2020). We used the Dsuite ‘Dtrios’ program to examine gene flow, using the same population partitions used in the HyDe analysis above. We also included the option to use a phylogenetic tree to specify the relationships between populations (see Supplementary Data tree.nwk). Full details are described under Step 2 of ‘Introgression’ at protocols.io under dx.doi.org/10.17504/protocols.io.bq7tmznn

#### Allele sharing

In order to quantify the extent of introgression in each genome we identified sites at which all samples of domestic ferrets where homozygous to a common allele and in which European polecats from the European mainland where homozygous to a common alternative allele at the same site (using GATK SelectVariants). For this we used all eight domestic ferret genomes (all of which had a minimum read coverage of 18.9x) and compared them to eight high-coverage (>10x) European mainland polecats. One sample (euro_LIB21977) was later removed from the analyses due to the suggestion of a small amount of genome introgression present in this sample (Figure *5*). We restricted our analysis to biallelic SNP sites and then summarised our results (using GATK VariantsToTable) to confirm that all the homozygous European polecat sites were alternate to the same allele. Next, we calculated the mean SNP quality score (QUAL) for each site and filtered out any sites with a score less than 40. We then took each British polecat sample in turn, identified the genotype at each corresponding site and calculated the proportion of the genome that had ferret-specific or polecat-specific alleles. Using the ONS shapefile for NUTS2 (https://geoportal.statistics.gov.uk/datasets/48b6b85bb7ea43699ee85f4ecd12fd36_0) we allocated each sample to one of the 40 different NUTS2 geographical regions and calculated the mean contribution of ferret-specific and polecat-specific alleles to polecat genomes in that region.

To calculate the false discovery rate (FDR) of this method, we took the 12 Mb capture array for the 76 domestic ferret samples (see Methods: Samples and Sequencing) and calculated genotypes at each of the fixed positions. We then identified the accuracy of our method by calculating the number of genotype calls in the 76 samples that agreed/disagreed with our predictions. Full details of the allele sharing protocol and FDR analyses are described under Step 3 of ‘Introgression’ at protocols.io under dx.doi.org/10.17504/protocols.io.bq7tmznn

#### Topology weighting analysis

Bi-allelic SNPs present in all British and mainland European polecats, domestic ferrets, and weasel with less than 5% missing taxa were extracted using bcftools (Danecek & McCarthy, 2017), and then phased and imputed using beagle v4.1 with a window size of 10,000bp and overlap of 1,000bp (Browning & Browning, 2007). Phylogenetic trees were then inferred across the longest 23 scaffolds (all at least 25Mb in length) as well as any where there is putative evidence of adaptive introgression. IQTREE v1.6.12 (Nguyen, Schmidt, von Haeseler, & Minh, 2015) was used to infer trees for each 50bp window (with at least 40 sites per individual), with an overlap of 10bp, using scripts adapted from genomics_general (https://github.com/simonhmartin/genomics_general). Automatic model selection (Kalyaanamoorthy, Minh, Wong, von Haeseler, & Jermiin, 2017) was carried out for each window with ascertainment-bias correction. Topology weighting analysis was then carried out using TWISST (Martin & Van Belleghem, 2017). The weightings were plotted in R (v3.5.2), after smoothing with a loess span of 0.05.

#### Adaptive introgression

In order to identify putatively introgressed regions in English polecats, we took the most likely trio of populations showing introgression (identified from our previous analyses) and filtered the input for scaffolds longer than 1Mb where all samples had called genotypes. We then used Dsuite Dinvestigate to identify sliding windows of 1000 SNPS and their associated admixture statistics (D,*f*_d_,*f*_dM_, and *df*) and selected the windows with the top 1-percent of *f*_dM_ values (Malinsky et al., 2020). The *f*_dM_ statistic (a modified version of the *f*_d_ statistic) is symmetrically distributed around zero under the null hypothesis of no introgression and can equally quantify shared variation between P3 and P2 (positive values) or between P3 and P1 (negative values).

Using VCFTools, we then identified the location of the top 1-percentile Fst outliers between the two ‘parental’ populations of the introgressed trio (Danecek et al., 2011). We filtered down the Fst outliers to those that overlapped with the 1-percentile *f*_dM_ windows and then identified which of these intersections overlapped with genes in the domestic ferret genome annotation (Peng et al., 2014)

## Results

### Phylogenetic analyses

We used RAxML to create ML phylogenies for the whole mitochondrial genomes (Figure *2*) and genome-wide SNPs (Figure *3*) (Stamatakis, 2014). In both phylogenies Steppe polecats and Black-footed ferrets form distinct clades, with Steppe polecat forming a sister clade to the European polecats. Also in both phylogenies, European mainland polecats form a paraphyletic group and there is a separate clade for domestic ferrets and British polecats and hybrids, the structure of which varies between the two phylogenies.

**Figure 2.**
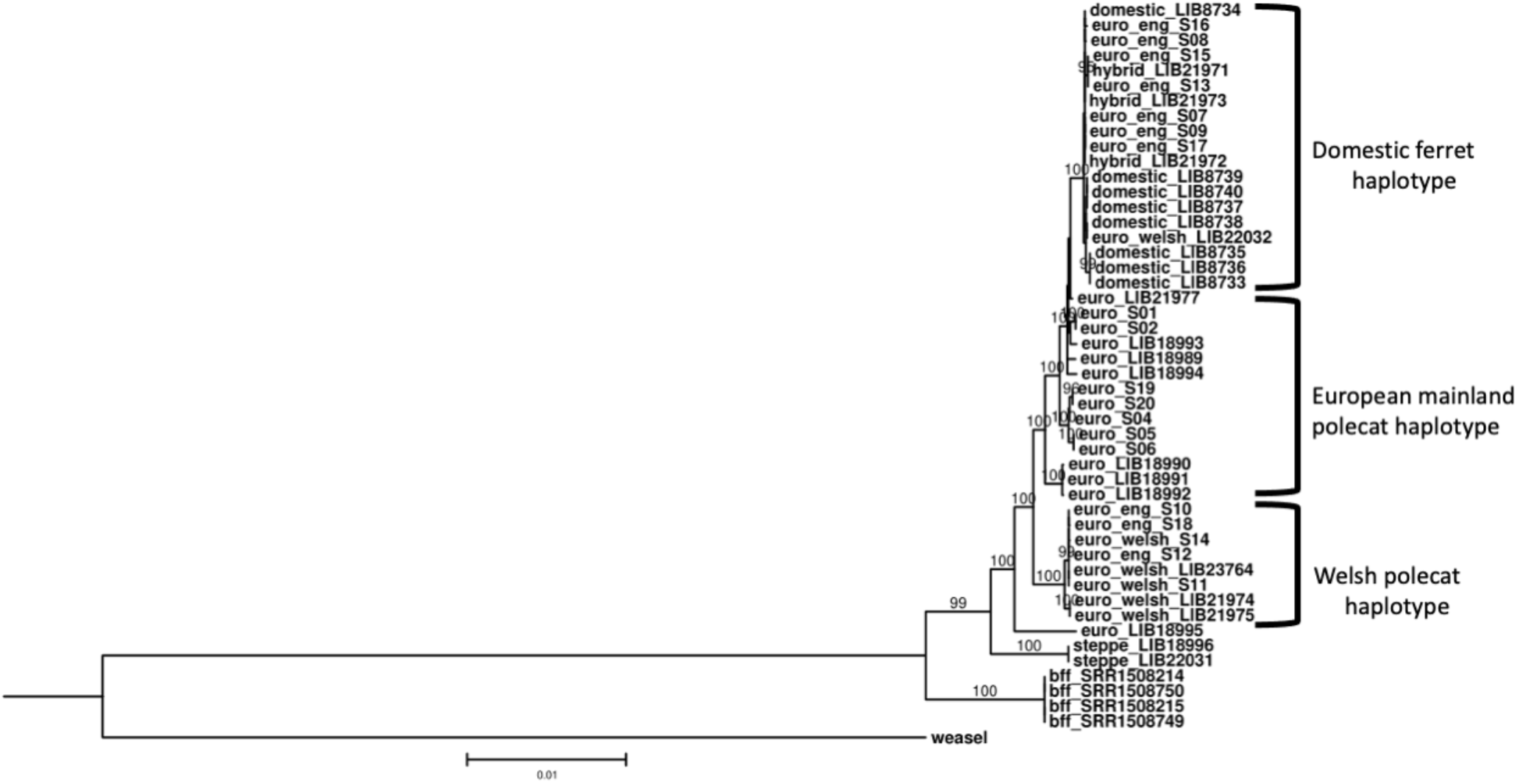
Maximum likelihood phylogeny for mitochondrial whole genomes with main haplotype groups annotated. Numbers at nodes refer to bootstrap support values of 95 and above. Domestic ferrets are labelled as ‘domestic’, European polecats from mainland Europe are labelled as ‘euro’, European polecats from England are labelled as ‘euro_eng’, European polecats from Wales are labelled as ‘euro_welsh’, and samples identified (by phenotype) as ferret x polecat hybrids are labelled as ‘hybrid’. Steppe polecats and Black-footed ferrets are labelled as ‘steppe’ and ‘bff’ respectively. The tree is rooted with Least Weasel and branch lengths are in expected substitutions per site.

**Figure 3.**
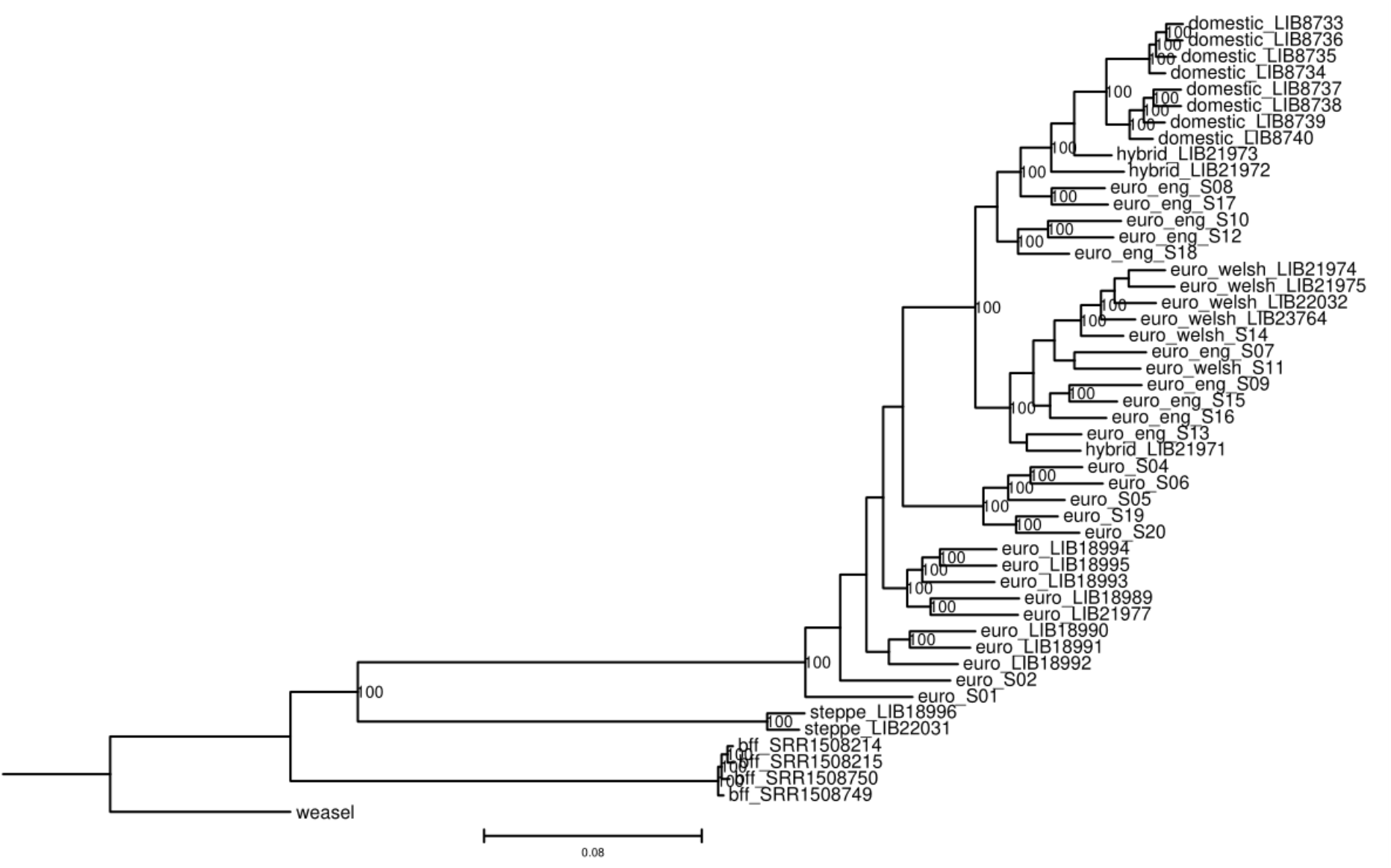
Maximum likelihood phylogeny for genome-wide SNPs. Numbers at nodes refer to bootstrap support values of 95 and above. Domestic ferrets are labelled as ‘domestic’, European polecats from mainland Europe are labelled as ‘euro’, European polecats from England are labelled as ‘euro_eng’, European polecats from Wales are labelled as ‘euro_welsh’, and samples identified (by phenotype) as ferret x polecat hybrids are labelled as ‘hybrid’. Steppe polecats and Black-footed ferrets are labelled as ‘steppe’ and ‘bff respectively. The tree is rooted with Least Weasel and branch lengths are in expected substitutions per site.

In the mitochondrial phylogeny (Figure *2*), all but one of the Welsh polecats form a clade with three English polecats (all from western England), with the remaining British polecats, domestic ferrets, and phenotypic hybrids forming a separate clade. This suggests two distinct mitochondrial haplotypes within Great Britain; one a Welsh polecat haplotype, unique from that found in mainland Europe, the other a domestic ferret haplotype. The multiple sequence alignment showed 78 out of 15511 sites segregated all samples of the Welsh polecat haplotype from all samples of the domestic ferret haplotype.

In the genome-wide SNP phylogeny (Figure *3*), British polecats are separated into two distinct clades – one containing domestic ferrets, two of the three hybrids, and five English polecats, the other containing all the Welsh polecats, along with five English polecats and the remaining hybrid. This clade also separates into two sub-clades, one containing all the Welsh polecats (with the addition of one English polecat, S07) and the other containing all remaining English polecats and hybrids. The three samples of English polecats that cluster with the Welsh polecats in mitochondrial tree (samples S10, S12, and S18) are all found with the domestic ferret branch of the genome-wide SNP tree but form a separate clade at the base.

#### European mainland polecats

Polecats from the European mainland also cluster by geographical region with samples originating from Austria, Germany, Italy, France, and Spain. (Supplementary Table S1). In the mitochondrial genome tree (Figure *2*), polecats from Austria and Italy cluster into well-defined clades, whilst the remaining samples from Germany, France, and Spain are less well defined. The strongest sub-population structure is observed in the genome-wide SNP phylogeny (Figure *3*). Samples from Austria and Italy form separate clades, along with a clade with samples from Spain and France, which are then separated into separate sub-clades of Spanish and French samples. Finally, the samples from Germany form nested distinct lineages at the base of the European polecat samples.

We used TreeMix to examine population splits and migration events (Pickrell & Pritchard, 2012), and calculated that the optimum number of migration edges as 2 (Figure *4*, Supplementary Figure S1). As is seen in the nuclear ML phylogeny (Figure *4*), Steppe polecat and Black-footed ferret form sister species, with mainland European polecat forming a single clade and British polecats and ferrets forming another clade. There is a strong migration edge from domestic ferret (‘domestic’) to English polecat (‘euro_eng’), suggesting gene flow between them, along with another migration edge (of a smaller migration weight) from Steppe polecat (‘steppe’) to mainland European polecat (‘euro’).

**Figure 4.**
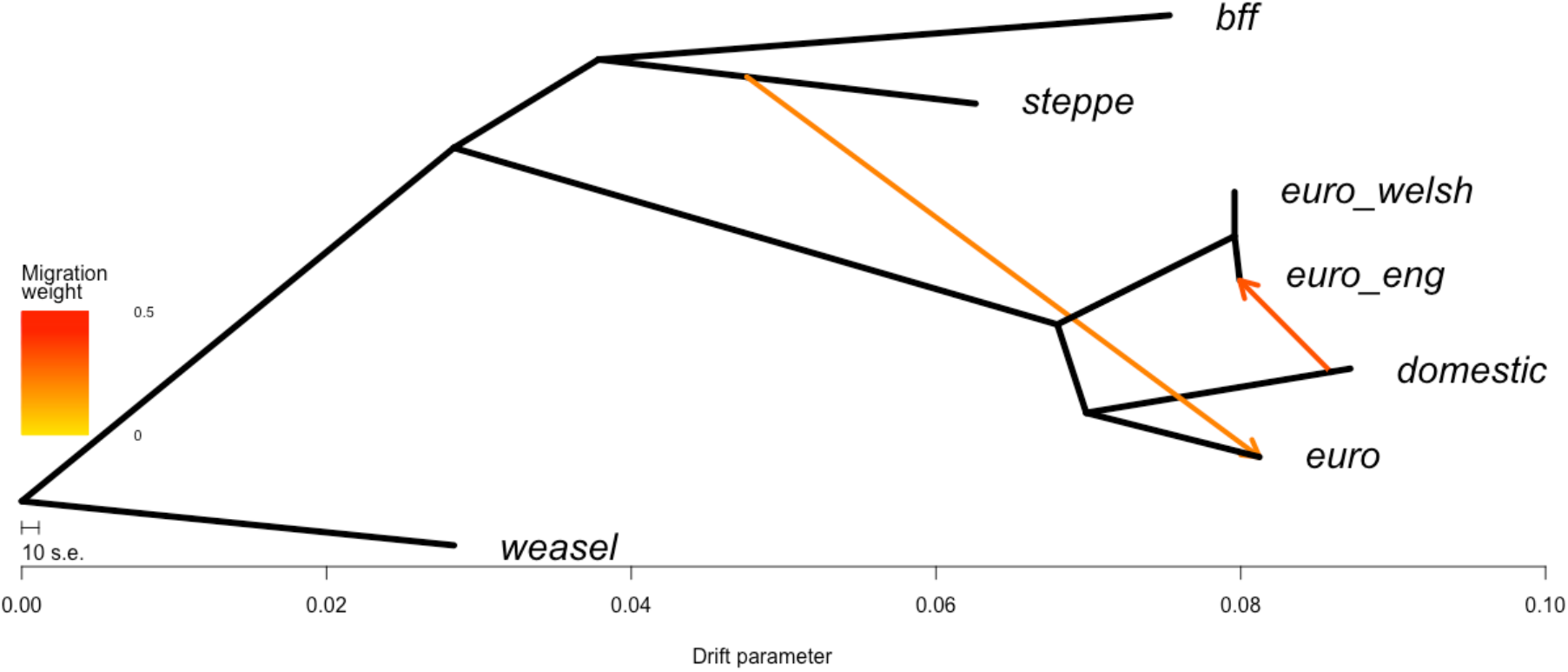
TreeMix phylogeny showing the optimum number of migration edges (two). The tree is rooted by Least Weasel (‘weasel’). Populations are labelled as in Figure 2.

### Population Structure

We used Admixture to plot population structure across 5 values of k (1-5) (Figure *5*) and calculated the CV error for each value of K (Supplementary Figure S2). CV-error increased noticeably after K=3. We included phenotypic polecat x ferret hybrids in the plots as a separate group to demonstrate the differences (and similarities) to those individuals that had been phenotyped as ‘pure’ European polecats. At least one of the hybrids has more polecat-like genetic structure (0.729) than some of the phenotypically pure English polecats (0.709 - 0.999), most of which show ferret-like genetic structure to a varying degree (Supplementary Table S3). Additionally, there is evidence of ferret introgression in one sample of European mainland polecats from Spain. Also, there is a suggestion of a unique genetic structure in the Italian population of polecats, albeit at K values (K=4 and K=5) higher than the most significant (K=3).

**Figure 5.**
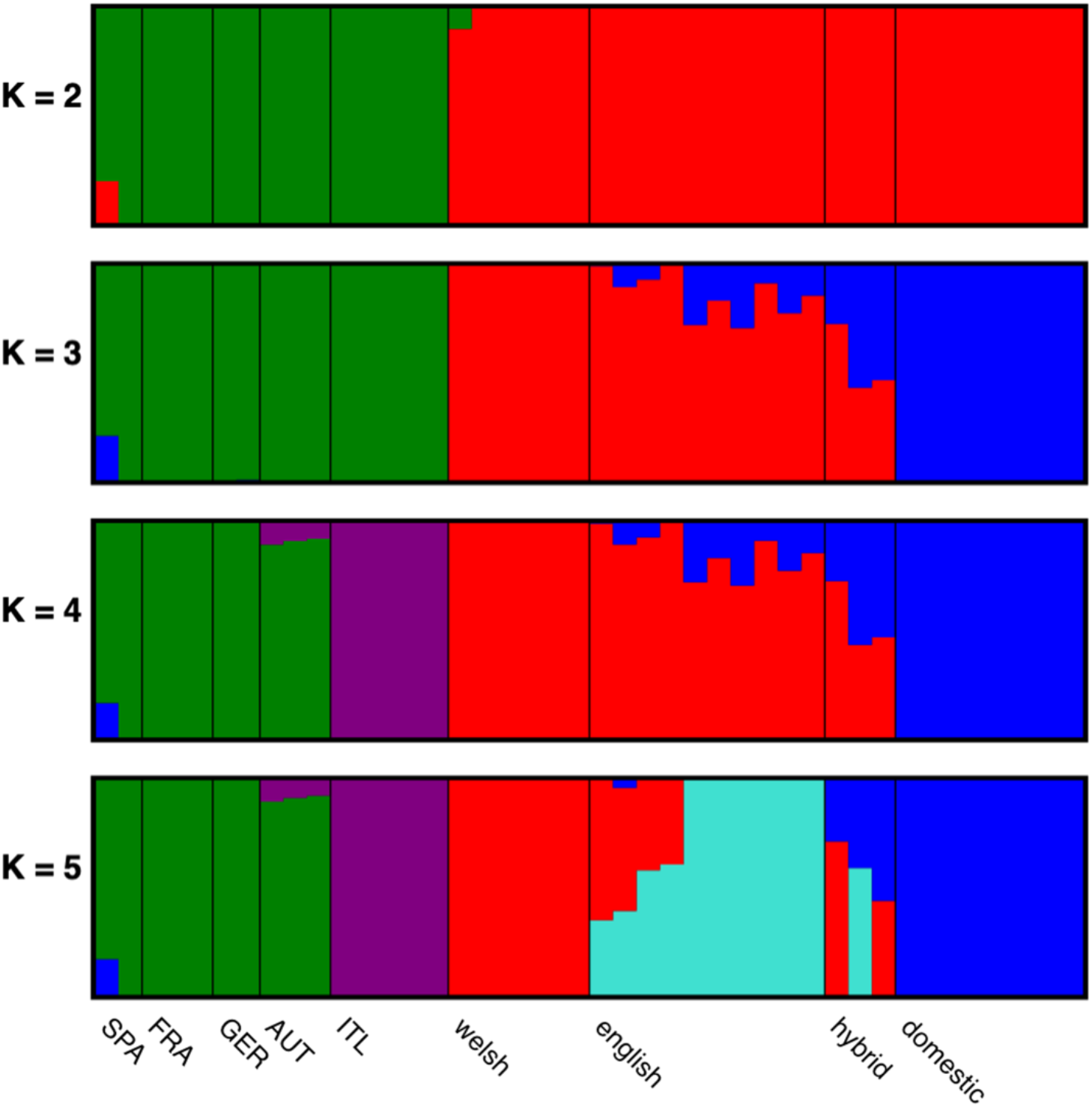
Admixture plots for values of K=2 (top) to K=5 (bottom). Samples from the European mainland have been further broken down into their country of origin as follows. SPA=Spain, FRA=France, GER=Germany, AUT=Austria, and ITL=Italy.

#### Introgression analyses

The significant results from Hyde identified four trios that showed evidence of hybridisation. Three out of the four trios identify English polecats as hybrids, with the top hit placing domestic ferrets as P1 and Welsh polecats as P2 (Table 1 and Supplementary Table S4).

**Table 1.** Filtered results from HyDe, ordered by p-value, then Z-score. ‘Gamma’ is an indication as to the proportion of genetic loci contributing from P1 and P2, where a value of 0.5 would indicate a 50:50 hybrid.

| P1 | Hybrid | P2 | Z-score | P-value | Gamma |
| --- | --- | --- | --- | --- | --- |
| domestic | euro_eng | welsh | 159.040951 | 0 | 0.33624814 |
| euro | domestic | welsh | 28.7314146 | 0 | 0.08208683 |
| euro | euro_eng | welsh | 38.0325902 | 0 | 0.05330685 |
| domestic | euro_eng | euro | 4.96293934 | 3.48E-07 | 0.98900496 |

We then tested for hybridisation at the individual level within the populations that have significant levels of hybridization (Supplementary Table S5).

Table 2 shows individuals across the hybrid population of the four trios listed in Table 1 that have a Gamma value > 0.2 and < 0.8. The only individuals that met this requirement were all the English polecats where P1 and P2 were domestic ferret and Welsh polecats respectively. Additionally, we noted that despite being phenotyped as a hybrid, individual hybrid_LIB21971 shows less evidence of hybridisation than seven English polecats which were all phenotyped as ‘pure’ polecats.

**Table 2.** Statistically significant HyDe tests for hybridisation on individuals of hybrid populations flagged as significant in Table 1 above and that have a Gamma value > 0.2 and < 0.8

| P1 | Hybrid | P2 | Zscore | Pvalue | Gamma |
| --- | --- | --- | --- | --- | --- |
| domestic | hybrid_LIB21973 | welsh | 200.10 | 0 | 0.533688723 |
| domestic | hybrid_LIB21972 | welsh | 191.14 | 0 | 0.569239275 |
| domestic | eng_euro_S17 | welsh | 186.55 | 0 | 0.422036059 |
| domestic | eng_euro_S10 | welsh | 149.59 | 0 | 0.372189884 |
| domestic | eng_euro_S18 | welsh | 158.44 | 0 | 0.350551109 |
| domestic | eng_euro_S08 | welsh | 164.65 | 0 | 0.346368603 |
| domestic | eng_euro_S12 | welsh | 129.86 | 0 | 0.298052044 |
| domestic | eng_euro_S07 | welsh | 130.62 | 0 | 0.275299731 |
| domestic | eng_euro_S09 | welsh | 136.08 | 0 | 0.266102112 |
| domestic | hybrid_LIB21971 | welsh | 133.00 | 0 | 0.256771089 |
| domestic | eng_euro_S13 | welsh | 123.43 | 0 | 0.234519286 |
| domestic | eng_euro_S16 | welsh | 116.13 | 0 | 0.231423632 |
| domestic | eng_euro_S15 | welsh | 104.01 | 0 | 0.208069401 |

To examine introgression in the context of gene flow between populations, we used Dsuite to calculate the D-statistic for the same combination of trios used in the HyDe analyses above (Malinsky et al., 2020). The output from *Dsuite Dtrios* echoed that of the filtered HyDe results in that the trio with the highest probability of gene flow were domestic ferret, English polecats, and European polecats (Table *3*). The highest-ranking score in this analysis placed polecats from the European mainland as the parental population, with Welsh polecats as the second-highest ranking result. Geographically, it would be impossible for polecats from the European mainland to be the parental population and this is probably a reflection of the similarity between polecats from Wales and the European mainland.

**Table 3.** Output of Dsuite Dtrios. The combination of P1, P2, and P3 are ordered so as the D statistic is always positive and all the results, including the f4-ratio reflect evidence of excess allele sharing between P3 and P2 for each trio.

| P1 | P2 | P3 | D-statistic | Z-score | p-value | f4-ratio |
| --- | --- | --- | --- | --- | --- | --- |
| euro | english | domestic | 0.322879 | 27.8886 | 0 | 2.01083 |
| welsh | english | domestic | 0.18104 | 20.112 | 0 | 1.57156 |
| euro | welsh | domestic | 0.196827 | 16.9025 | 0 | 1.26856 |
| welsh | english | euro | 0.0345665 | 6.92304 | 2.21E-12 | 0.12141 |

We identified 15,755 sites that were homozygous reference in 8 high-coverage ferret genomes (ferret-specific alleles) and homozygous alt (all to the same alt allele) in the 7 high-coverage European mainland polecats (polecat-specific alleles). We calculated the number of ferret-specific and polecat-specific alleles in each British polecat sample and mapped the distribution of ferret-specific and polecat-specific alleles across Great Britain, along with the sample-specific mitochondrial haplotypes, assigned in Figure *2* (Figure *6*).

**Figure 6.**
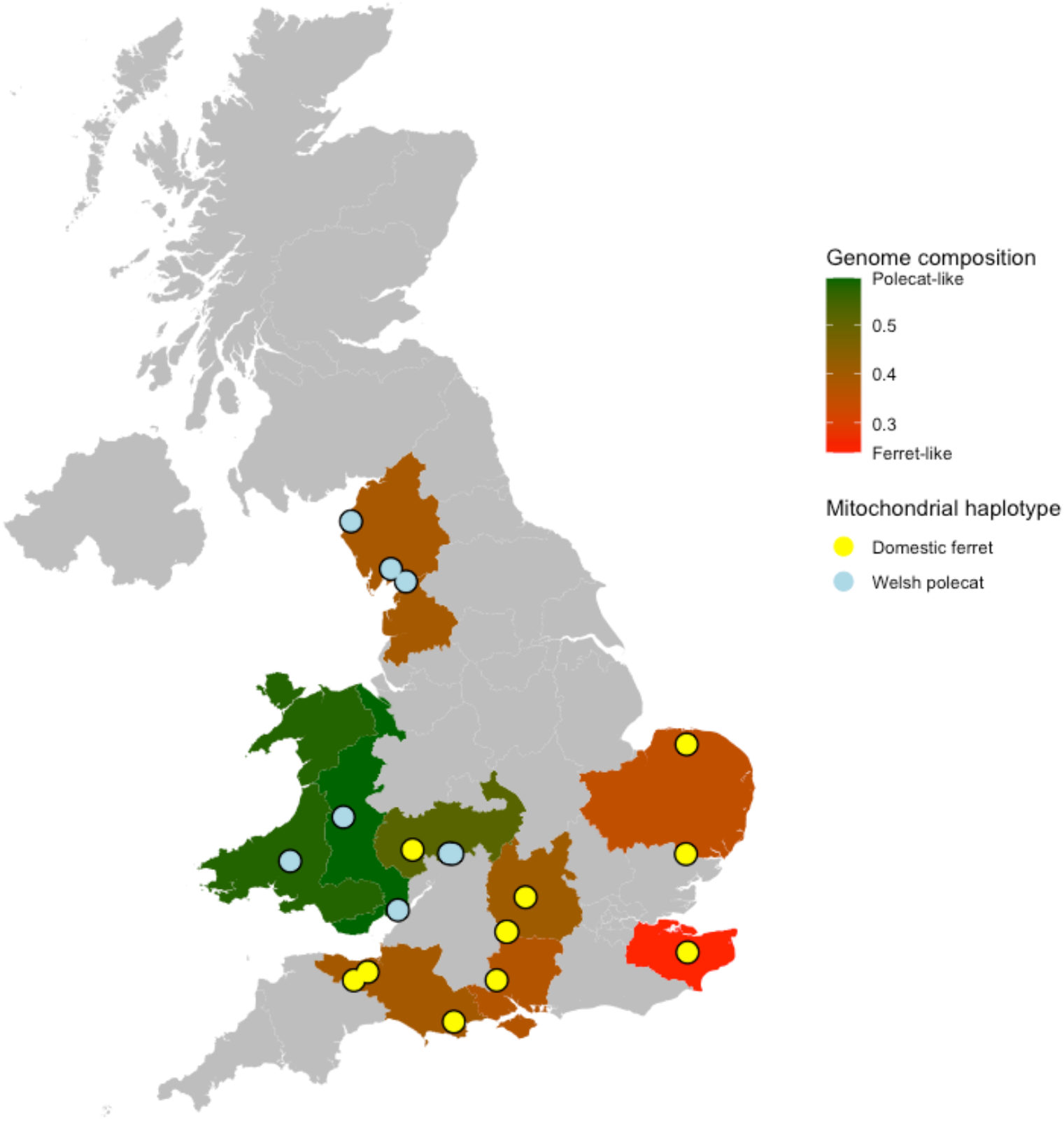
Distribution of polecat-specific and ferret-specific alleles across Great Britain, along with mitochondrial genome haplotypes assigned in Figure 2A.

As can be seen as the distance increases away from the Welsh refugia, polecats become increasingly ferret-like in their genome composition, with an individual in the south-east of England being the most ferret-like. Regions where polecats persisted during the 1900s bottleneck (Wales and English counties bordering Wales) and regions where Welsh polecats were reintroduced (Cumbria), still show predominantly polecat mitochondrial haplotypes (the exception being sample LIB22032), whilst all other samples have ferret mitochondrial haplotypes producing a clear division in haplotypes. Polecats in Wales show the greatest genetic identity to those from mainland Europe, followed by central western England and Cumbria.

Finally, we calculated the FDR of this method on a 12 Mb capture array across 76 domestic ferret samples. Across the 76 samples 2,382 genotype were called, 2,362 of which were called as homozygous ref, giving an FDR of 0.0008.

#### Topology weighting analysis

The topology weightings out of the fifteen possible topologies of the five populations (weasel, Welsh polecats, English polecats, mainland European polecats and domestic ferrets) were compared to the genome (Figure *7*). The most frequent topology shows the domestic ferret as most closely related to the English polecats. The next most frequent shows the English and Welsh polecats as sister to each other, with the domestic ferret most closely related to this group (Figure *7* A). The genome-wide distribution of the latter topology shows a relatively equal range of distribution across the genome (Figure *7* A and B (middle panel in blue)), whereas the first topology shows distinct peaks in super-scaffolds 73 and 139 (Figure *7* B; upper panel in green). A topology similar to the treemix-inferred one (Figure *4*), where the domestic ferret is closest related to the European mainland polecat, is only the fourth most frequent topology (Figure *7* A), and is fairly uniform across the genome, with a few distinct short spikes (e.g. Super scaffold 12, 46, 73).

**Figure 7.**
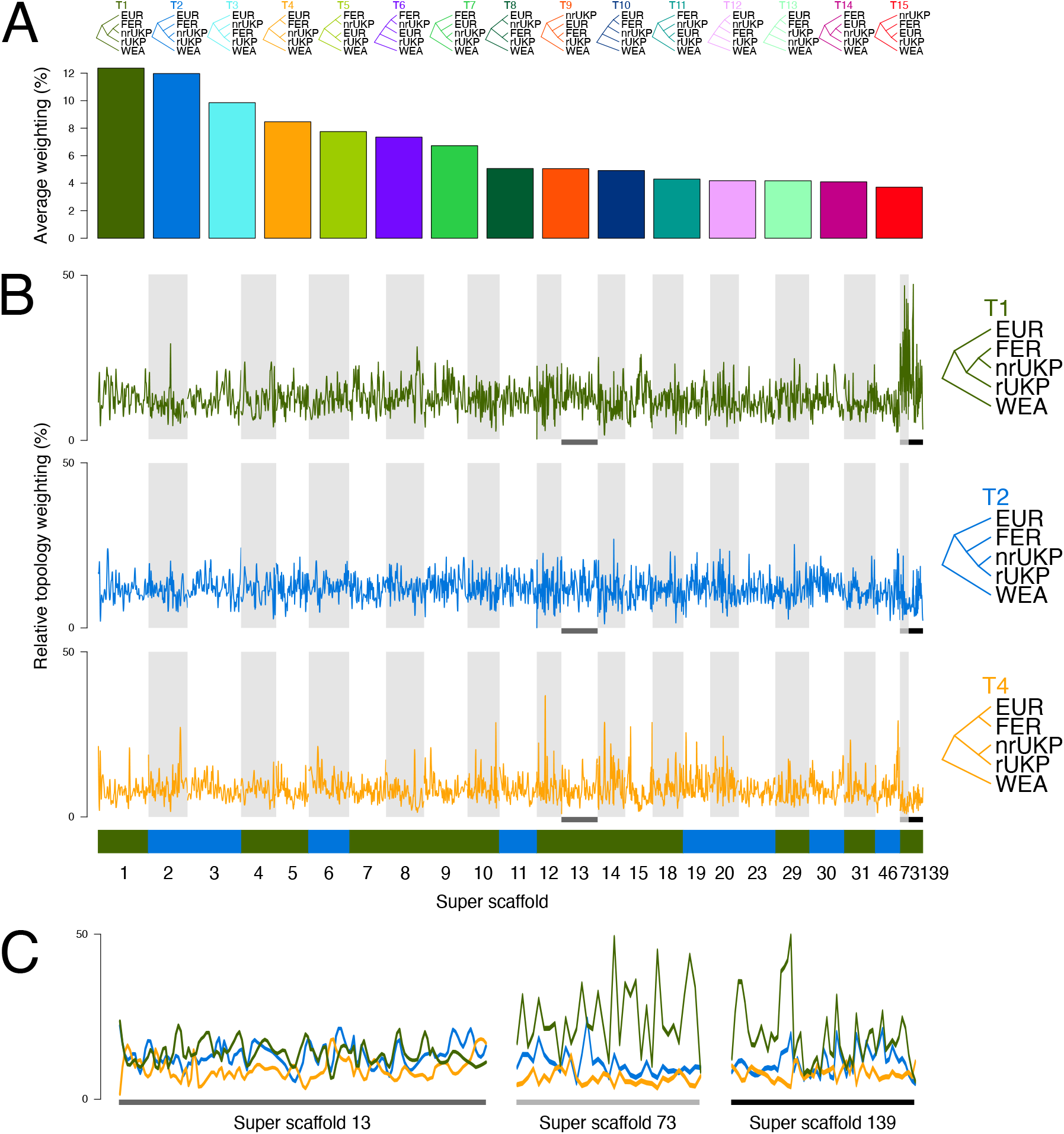
TWISST analyses of domestic ferret and polecat populations along the 23 longest scaffolds, and scaffolds associated with putative adaptive introgression (A) shows the genome-wide average weighting of the 15 possible topologies with the 5 specified population. B) shows the weighting of the two most frequent topologies, as well as the topology indicated by treemix analysis, across the scaffolds. The colour blocks below the panels show which topology out of the 15 was the most frequent overall across each scaffold, using the colours shown in panel A (all are either T1 or T2). C) shows the three super scaffolds putatively associated with adaptive introgression in more detail, with all three topologies from B plotted. The grey or black x axis bars correspond to the highlighted sections in B. The y axis in C is the same as B. Abbreviations: WEA – weasel; EUR – mainland European polecat; FER – domestic ferret; nrUKP – English polecat; rUKP – Welsh polecat.

### Adaptive Introgression

From our analyses above, it is apparent that English polecats show introgression from hybridisation between domestic ferrets and Welsh or European mainland polecats. Given the geographic limitations of our populations, we investigated adaptive introgression in English polecats (P2), using Welsh polecats (P1) and domestic ferrets (P3) as the two parental populations. We selected the top 1-percentile of windows that had positive *f*_dM_ values (51 windows) and identified 148 Fst outliers, all of which overlapped 7 of the 51 introgressed windows. The 7 windows spanned three scaffolds and composed either a single window within a scaffold or multiple overlapping windows forming a single continuous window in that scaffold (i.e. all three scaffolds contained only one continuous introgressed region) (Table *4* and Figure *7* C). Across the three scaffolds, these windows contained two proteincoding genes in the domestic ferret genome (ENSMPUG00000001059 and ENSMPUG00000008056), both of which are associated with cognitive function and sight in humans.

**Table 4.** Introgression and Fst data showing the number of introgressed windows in the top 1-percentile found in each scaffold, along with the combined span of the windows, the number of Fst ouliers found within those windows, and the number of protein-coding genes that intersected both the windows and Fst outliers.

| Scaffold name | N. of introgressed windows | Total span of introgressed windows | Fst outliers | N. protein coding genes |
| --- | --- | --- | --- | --- |
| Super-Scaffold 13 | 1 | 309Kb | 140 | 1 |
| Super-Scaffold 73 | 3 | 4.2Mb | 4 | 1 |
| Super-Scaffold 139 | 3 | 1.6Mb | 4 | 0 |

Of the total 148 Fst outliers, 140 of them occur within one 309Kb window on Super-Scaffold 13 (Table 4 and Figure 8). Within this region there is one protein-coding gene, ENSMPUG00000008056. The human ortholog of this gene, C19orf12, encodes for a mitochondrial associate transmembrane glycine zipper that is thought to act as a regulatory protein for magnesium transporters. Mutations in this gene are associated with a range of conditions, such as neurodegeneration with brain iron accumulation (NBIA), progressive movement disorders, spasticity, neuropathy, cognitive dysfunction, and optic nerve atrophy (Dusek et al., 2020; Espinos et al., 2020; Hartig et al., 2011; Landoure et al., 2013; Venco et al., 2015).

**Figure 8.**
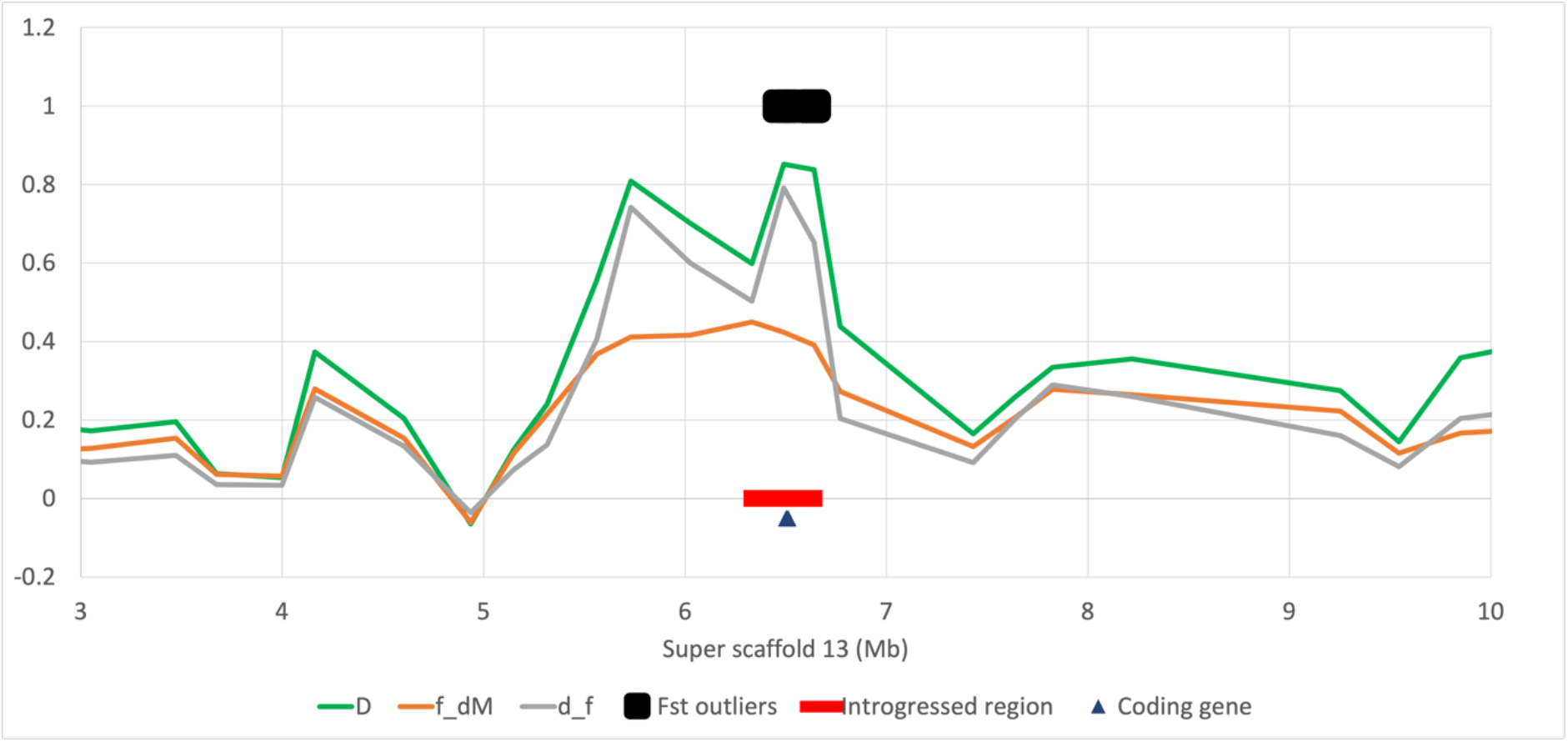
Admixture statistics for f_d_, f_dM_, and d_f_ across a 7Mb stretch of Super-scaffold_13. The red bar highlights the window of 1000 SNPs in the top 1-percentile of f_dM_ values, the black rectangle represents a cluster of 140 Fst outliers, and the black triangle represent the location of the protein-coding gene ENSMPUG00000008056.

The other coding gene, ENSMPUG00000001059, found on Super-Scaffold_73 (Supplementary Figure S3), is orthologous to the human CNTNAP5 gene, one of the contactin-associated proteins (Caspr), that participate in nerve excitation and conduction, along with neurotransmitter release in myelinated axons. Mutations in CNTNAP5 are associated with autism and neurodevelopmental disorders (Aleo et al., 2020; Ludington, Yu, Bae, & Barnett, 2020; Narita et al., 2020) as well as glaucomatous neurodegeneration (Chakraborty et al., 2021),

Finally, although no protein coding genes intersect with the Fst outliers on Super-Scaffold_139 it should be noted that ENSMPUG00000010497, orthologous to the Human RP1 axonemal microtubule associated gene, is located less than 15Kb downstream from the Fst ouliers. RP1 is a key gene in the formation of the outer segment of rod and cone photoreceptors in the eye (Goldberg, Moritz, & Williams, 2016). Mutations in RP1 are a common cause of retinitis pigmentosa, which involves a breakdown and loss of cells in the retina (Liu, Zuo, & Pierce, 2004).

## Discussion

Hybridisation and introgression may increase the adaptive potential of species by the fixation of beneficial alleles or removal of detrimental alleles and can be used as a tool for species recovery (Chan, Hoffmann, & van Oppen, 2019; Quilodran, Montoya-Burgos, & Currat, 2020). Whole genome sequencing has allowed quantitative assessment of introgression in British polecats. Our analyses show that British polecats, away from the previous refugia of central Wales show varying degrees of introgression with domestic ferrets. Our nuclear phylogeny (using genome wide SNPs) illustrates two clades in British polecats, one where some English polecats cluster with domestic ferrets, the other where the remaining English polecats cluster with Welsh polecats (Figure *3*). There are two well-supported mitochondrial haplotypes in British polecats – a polecat-like haplotype, confined to Cumbria, Wales and English counties bordering Wales, and a domestic ferret-like haplotype, found across the remaining range (Figure *2*). Previous work suggested the presence of two mitochondrial haplotypes in British polecats, one found in Cumbrian polecats, the other in Welsh polecats (Costa et al., 2013). We found no evidence to support this, with samples from both areas occurring within the same clades and no segregating SNPs between samples from the two proposed groups, supporting that found by (Davison et al., 1999) (although (Costa et al., 2013) had a greater number of samples than in our study (169 samples classified as polecat or hybrid)). The placements of British polecats across the two mitochondrial clades are inconsistent with those in the nuclear phylogeny, suggesting a high degree of hybridisation and introgression in British polecats outside Wales. Additional evidence supporting introgression within British polecats can be seen in the TreeMix analysis, which allows both population splits and migration events, and shows a high migration weight between English polecats and domestic ferret (Figure *4*). ADMIXTURE analysis further demonstrates introgression between domestic ferrets and English polecats. Additionally, it shows that many of the English polecats phenotyped as ‘pure’ polecats, show close to, or in some cases, more introgression than those phenotyped as hybrids. Another striking feature of the ADMIXTURE analysis is that polecats from the European mainland are allocated a separate genetic cluster than those from Wales (Figure *5*). Taking this, along with the phylogenetic analyses, suggests that Welsh polecats may form a separate phylogenetic unit from those on mainland Europe, driven by historical geographical separation between the two populations. Further analyses using HyDe (Table 1 and Table 2) and Dsuite (Table 3) shows that the most likely trios for significant hybridisation and introgression are Welsh polecats, domestic ferrets, and English polecats, with English polecats being the introgressed population. Again, in the HyDe analyses an individual phenotyped as a hybrid appears less introgressed than several individuals phenotyped as pure polecats. Finally, six samples of English polecats (S07, S09, S13, S15, S16, and hybrid_LIB21971) consistently show lower levels of introgression than the remaining English polecats. They all cluster with the Welsh polecats in the genome-wide SNP phylogeny, have Gamma values of less than 3 in the individual Hyde analyses and show the lowest values of ferret genetic structure in the ADMIXTURE analysis.

We have visualised the distribution of polecat mitochondrial haplotypes and nuclear alleles across the British range of the polecat (Figure *6*). Polecat mitochondrial haplotypes are restricted to north and central western Britain. The further away from the 1900s central Wales refuge polecats occur, the more ferret-like their genomes become, consistent with previous work suggesting that male polecats expand their range away from the refuge and hybridise with female feral domestic ferrets (Davison et al., 1999).

Looking at the genome in more detail, the TWISST analyses show that considering English polecats as a separate population, most regions of their genomes show signs of introgression with domestic ferret, although small windows of genome with no introgression remain. The introgressed regions of the genomes of English polecats contain two (and possibly three) genes that are associated with cognitive function and sight in humans.

Finally, we uncover novel features for the polecats from the European mainland. Firstly, there is evidence of introgression between European polecats and Steppe polecats (Figure *4*), supporting previous phenotypic analysis where the two species occur in sympatry (Cserkész et al., 2021). Also, there is a suggestion of genetic structure in mainland Europe (Figure *5*) where polecats from Italy show a distinct genetic population structure when compared to that of other countries.

## Supporting information

tree.nwk

## Summary

For the first time, we have carried out population-level whole genome sequencing on European polecats. Our analyses show that English polecats are highly introgressed with domestic ferrets, showing geographically separated mitochondrial haplotypes, with polecats becoming increasingly introgressed with domestic ferrets as they move further away from the 1900s polecat refugia. Phenotyping polecats as ‘pure’ or ‘hybrid’ is not accurate, with all samples of English polecat, regardless of phenotype showing some degree of introgression, with those assigned as ‘hybrid’ sometimes being less introgressed than those assigned as ‘pure’. Introgression is distributed widely across the genome and genes under selection for the introgressed regions include those that may assist in cognitive function and sight.

## Data availability

The domestic ferret reference genome used in this study can be found under ENA accession number GCA_920103865. Read data for Black-footed ferrets can be found at NCBI under BioProject PRJNA254451. Read data for the remaining samples can be found under ENA Study accession PRJEB48359 and described in detail in Supplementary Table S6.

## Acknowledgements

We’d like to thank Katherine A. Sainsbury, The Vincent Wildlife Trust, and National Museums Scotland for access to British polecat samples, Angus Davison (University of Nottingham) for providing the Steppe polecat samples, and Claudine Montgelard (Centre for Functional and Evolutionary Ecology, Campus du CNRS) for providing the Least Weasel sample. We acknowledge funding from the Biotechnology and Biological Sciences Research Council (BBSRC), part of UK Research and Innovation, Core Capability Grant BB/CCG1720/1 and the work delivered via the Scientific Computing group, as well as support for the physical HPC infrastructure and data centre delivered via the NBI Computing infrastructure for Science (CiS) group. Next-generation sequencing was delivered via the BBSRC National Capability in Genomics and Single Cell (BB/CCG1720/1) at Earlham Institute by members of the Genomics Pipelines Group.

## Authors Contributions

GJE, EC, JM, WH, and FDP designed the research, GJE performed the research and analysed the data (AC performed and analysed the Topology Weighting analyses and RS performed and analysed the genome-wide SNP phylogeny), EC and JM contributed numerous samples, GJE and WH wrote the paper.

## Competing Interests

The authors declare no competing interests.

---

**Supplementary Figure S1.**
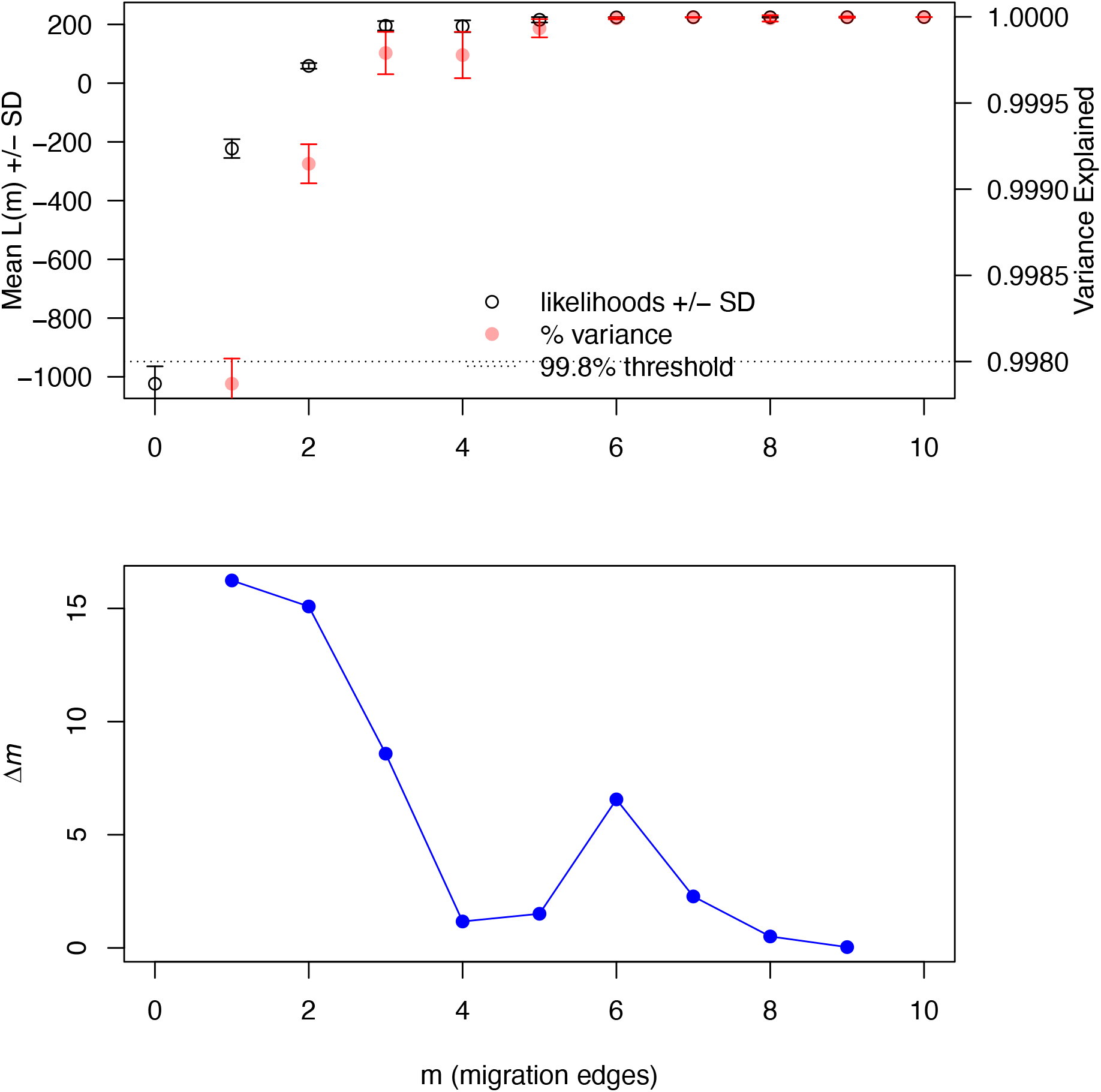
Estimation of the optimal number of migration edges using the R package OptM with the default Evanno method. The number of migration edges (2) was chosen based on a plateau of log likelihoods and when greater than 99.8% of the variance was explained.

**Supplementary Figure S2.**
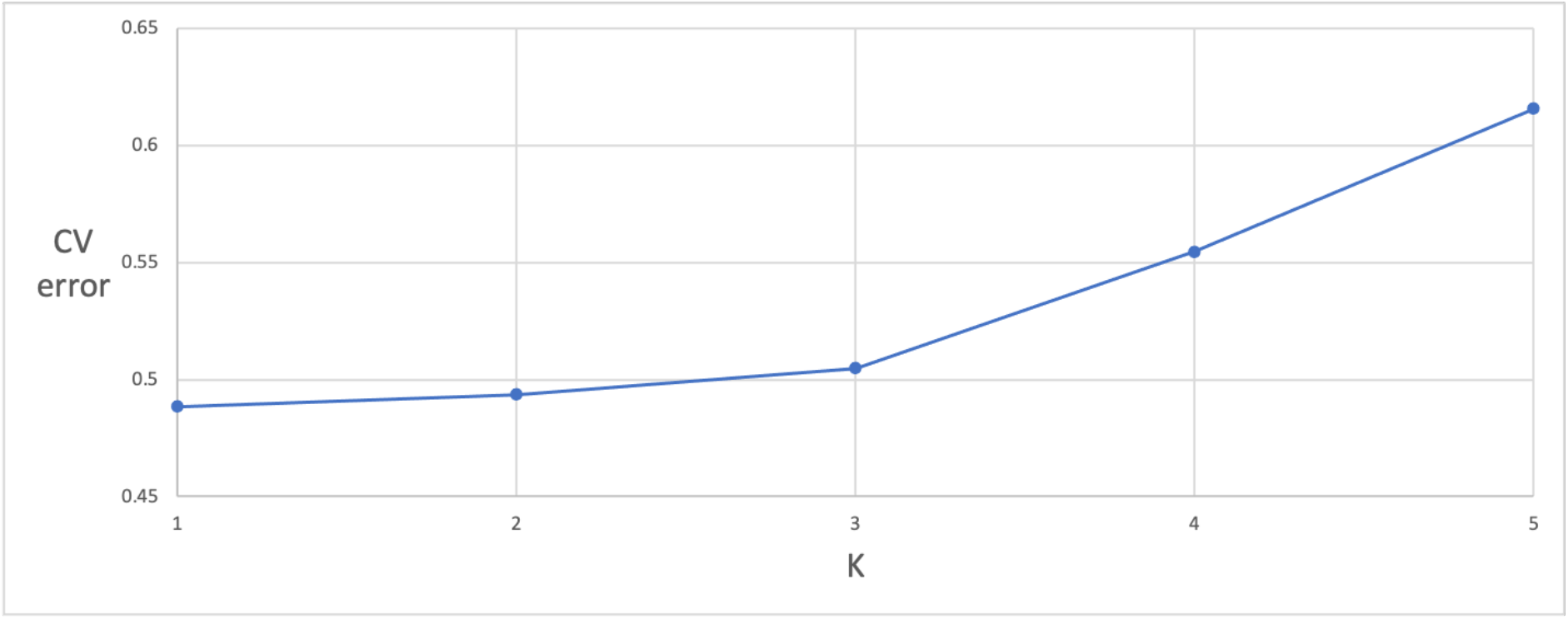
Calculation of CV error for each value of K in the Admixture analyses. CV-error increases noticeably after K=3.

**Supplementary Figure S3.**
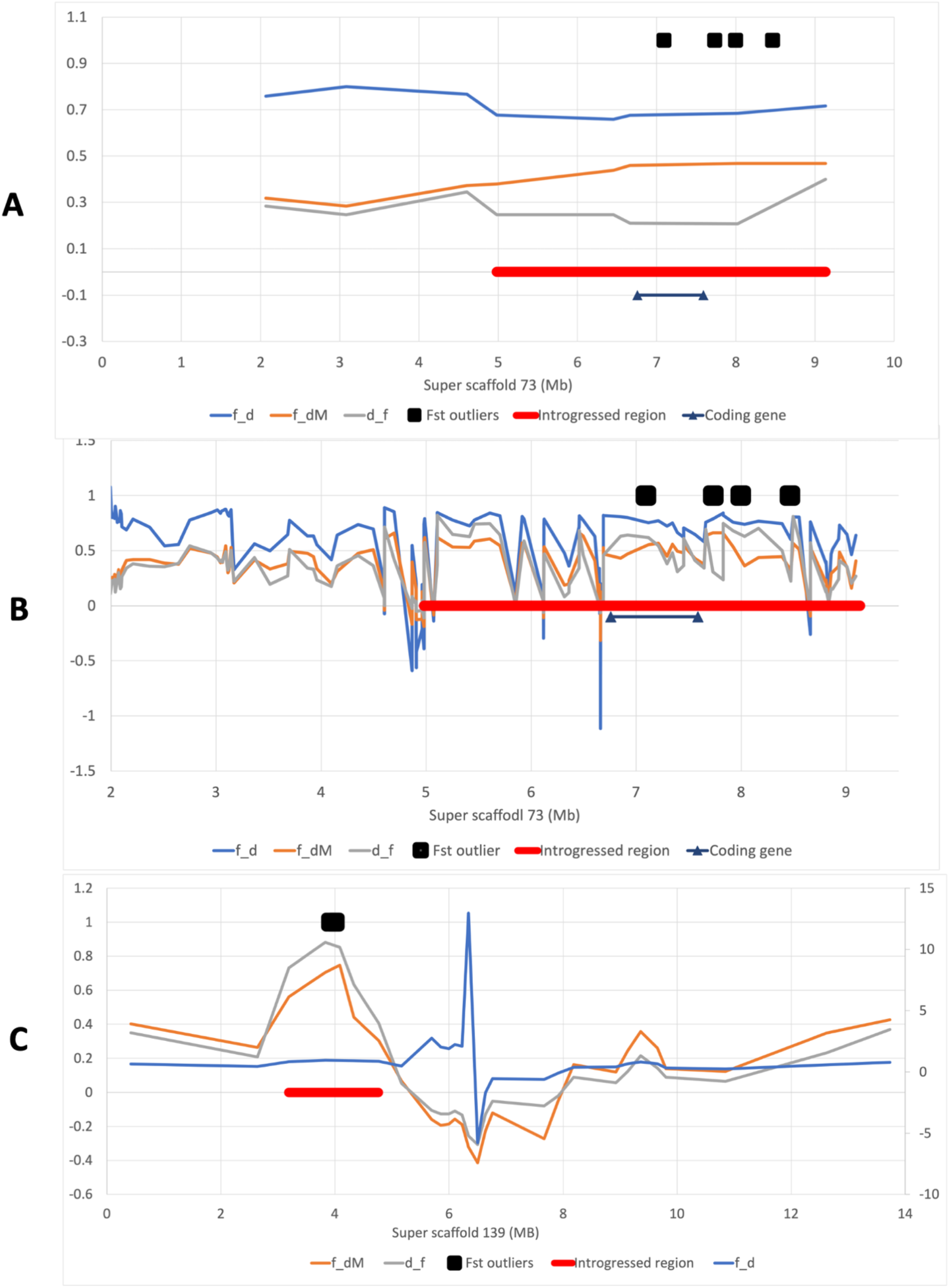
Admixture statistics for f_d_, f_dM_, and d_f_ across Super-scaffold_73 over A) Windows of 1000 SNPs, and B) windows of 50 SNPs (to view more local variation across the region), and C) Super-Scaffold_139. The red bar highlights the overlapping windows of 1000 SNPs in the top 1-percentile of fdM values, the black squares represent Fst outliers, and the area between the black triangles represent the location of the protein-coding genes.

## Other supplementary data (See supplementary files)

Supplementary Table S1. Details of per-sample information. ‘Sample Accession ID’ refers the ENA sample accession ID, and ‘Analyses population’ refers to the population (described in the manuscript) that the sample was allocated.

Supplementary Table S2. Sequencing information for all samples. Bases per sample and read coverage (based on the 2.42Gb domestic ferret genome) refers to post-QC data. Sequenced read-length refers the read-length pre-QC. Samples can be matched to those in Supplementary Table S1 by their Library ID.

Supplementary Table S3. Raw output of the Admixture program, providing genetic partitions for k=3.

Supplementary Table S4. Filtered output of ‘run_hyde.ph’. Columns containing only zeros have been removed.

Supplementary Table S5. Output of the ‘individual_hyde.ph’. Columns containing only zeros have been removed and an extra column ‘Gamma diff’ as been inserted to calculate the distance of Gamma (an indication as to the proportion of genetic loci contributing from P1 and P2, where a value of 0.5 would indicate a 50:50 hybrid), from 0.5. The table is sorted by p-value and then Z-score.

Supplementary Table S6. Full details of all ENA accession IDs for samples, experiments, and runs. Samples can be matched to those in Supplementary Tables S1 and S2 by their Library ID.

tree.nwk. Phylogenetic tree in Newick format, specify the relationships between populations in the Dsuite analyses.

## Notes

https://www.ebi.ac.uk/ena/browser/view/GCA_920103865

https://www.ebi.ac.uk/ena/browser/view/PRJEB48359

### Competing Interest Statement

The authors have declared no competing interest.

